# Connectome embedding in multidimensional graph-invariant spaces

**DOI:** 10.1101/2023.04.25.538320

**Authors:** Mathieu Mach, Enrico Amico, Raphaël Liégeois, Maria Giulia Preti, Alessandra Griffa, Dimitri Van De Ville, Mangor Pedersen

## Abstract

The topological organization of brain networks, or connectomes, can be quantified using graph theory. Here, we investigated brain networks in higher dimensional spaces defined by up to ten node-level graph theoretical invariants. Nodal invariants are intrinsic nodal properties which reflect the topological characteristics of the nodes with respect to the whole network, including segregation (e.g., clustering coefficient) and centrality (e.g., betweenness centrality) measures. Using 100 healthy unrelated subjects from the Human Connectome Project, we generated multiple types of connectomes (structural/functional networks and binary/weighted networks) and embedded the corresponding network nodes (brain regions) into multidimensional graph spaces defined by the invariants. First, we observed that nodal invariants are correlated between them (i.e., they carry similar network information) at a whole-brain and subnetwork level. Second, we conducted a machine learning analysis to test whether brain regions embedded in multidimensional graph spaces can be accurately classified into higher order (association, subcortical and cerebellar) and lower order (visual, somatomotor, attention) areas. Brain regions of higher order and lower order brain circuits were classified with an 80-87% accuracy in a 10-dimensional (10D) space. 10D graph metrics performed better than 2D and 3D graph metrics, and non-linear Gaussian kernels performed better than linear kernels. This suggests a non-linear brain network information gain in a high-dimensional graph space. Finally, we quantified the inter-subject Euclidean distance of each brain region embedded in the multidimensional graph space. The inter-individual distance was largest for regions of the default mode and frontoparietal networks, providing a new avenue for subject-specific network coordinates in a multidimensional space. To conclude, we propose a new framework for quantifying connectome features in multidimensional spaces defined by graph invariants, providing a new avenue for subject-specific network coordinates and inter-individual distance analyses.

## Introduction

Network theory has become an emerging avenue of investigation in science [1–6]. Network analysis is particularly relevant in neuroscience since the brain and its neurons comprise complex and multiscale interconnected networks [7–15]. A better comprehension of brain networks is a critical element in searching for simple and non-invasive diagnostic markers of neuropsychiatric and neurological diseases [16, 17], but also for the general understanding of how the different brain structures interact [18, 19]. An essential idea of network theory is the concept of node-level invariants, which are nodal scores reflecting the nodes’ ‘importance’ or topological role in the whole network. In graph-theoretical terms, invariants can be divided into integration measures (e.g., centralities) and segregation measures (e.g., clustering coefficient). Using graph invariants, we can also build a ranking of nodes and compare different nodes.

In computational and clinical neurosciences, many studies have used invariants at different scales to characterize the brain organization in healthy and diseased populations [13, 17, 24, 25], with popular invariant measures being the degree, betweenness, closeness, and eigenvector centrality [20–23]. Moreover, invariants can be combined and considered jointly within multidimensional spaces. For example, Joyce and colleagues [22] defined a new invariant called leverage centrality and compared it to three well-known invariants in connectomes (namely, degree, betweenness, and eigenvector centrality). To this end, they defined two-dimensional (2D) and three-dimensional (3D) spaces, where the different invariants represent the spaces’ dimensions and the single brain regions are points living in these spaces. Zuo and colleagues [23] used voxel-level invariant scatter plots to investigate the relationship between different invariants of brain functional connectomes. Nonetheless, most work on graph invariants applied to connectomes has been done in 1, 2, or 3D spaces, i.e., considering a few invariants at a time.

Our current study aims to extend these works by investigating brain connectomes within high-dimensional invariants’ spaces. To this end, we considered up to 10 graph invariants and used them to define multidimensional spaces within which we characterized inter-nodal and inter-subject connectomes’ distances. Our work proposes a new way to use graph invariants by creating Euclidean spaces where each axis represents a graph invariant. Each node (brain region) has a precise position in these spaces, which is defined by its invariant scores that make a set of coordinates.

Three main analyses were performed in this study. First, to understand similarities and dissimilarities between different invariants, we conducted a correlation analysis between invariants, both at whole-brain and functional subnetworks’ levels. Second, we explored the multidimensional graph spaces derived from up to 10 invariants using machine learning (ML), to test whether brain regions belonging to higher order and lower order brain circuits can be differentiated in such spaces. Finally, we characterized each brain region based on its inter-individual mean distances in multidimensional graph spaces. This analysis allowed us to identify the brain regions yielding the largest inter-individual variability in a multidimensional graph space.

## Results

Our approach comprises three steps and is schematized in Figure 1. First, the different invariants are computed from each connectome (Figure 1 a). The ten invariants considered in this work are listed and explained in more detail in the Methods, section 5.3. The rationale for investigating ten invariants is to understand which additional information about the human connectomes we can gain compared to considering smaller invariants’ subsets. It also allows exploring different combinations of invariants in lower dimensional spaces (2D and 3D, for example) to study connectomes. The next step is to interpret each computed invariant as an axis of a multidimensional Euclidean space, and the invariant scores of each brain region (network node) as coordinates in such space (Figure 1 b). This results in embedding each node in a new multidimensional space that we name ”graph space”. The value of this embedding is that brain regions are placed in the graph space according to their properties with respect to the whole-brain network and can be easily compared between each other and across different subjects. The final step is to explore this new space. We propose to compute pairwise distances between brain regions in the graph space. We used the Euclidean distance since we are working in a Euclidean space, and it can be very easily defined and calculated in a space of any dimension (Figure 1 c). The Euclidean distance can also be used to compare connectomes from different subjects at multiple scales (Figure 1 d)—for example, considering a single brain region across subjects or an average global distance.

**Figure 1:**
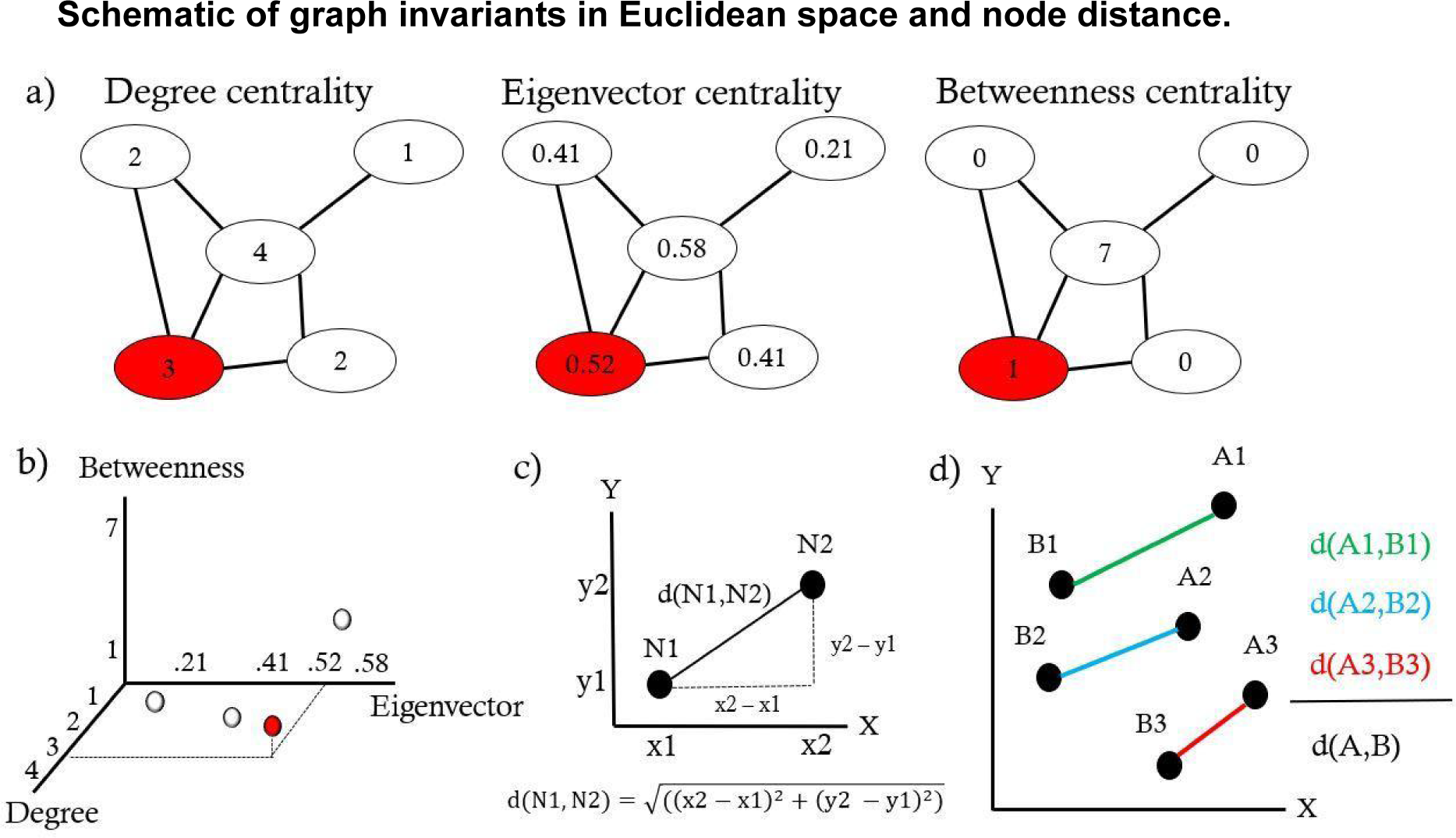
(a) Nodal degree, eigenvector, and betweenness centrality invariants computed on the same toy graph. (b) Illustration of the three-dimensional metric space (3D graph space) built from the above-listed invariants (degree, eigenvector and betweenness centrality). Each dot represents a node of the toy graph in (a). As an example, the red dot corresponds to the red node. (c) Euclidean distance *d* between two graph nodes embedded in a two-dimensional space. (d) Euclidean distances between homologous brain regions (regions 1, 2, and 3) from different subjects (A and B), and the average distance between the two subjects across homologous brain regions.

### Graph invariants are correlated at whole-brain and sub-network levels

We derived group-representative connectomes from structural, diffusion and resting-state functional magnetic resonance imaging (MRI) data of 100 healthy subjects of the Human Connectome Project [26], each one composed of 219 nodes (brain regions). The 219 regions were grouped into nine resting state networks (RSNs) [18]. Multiple connectome models were considered: binary structural and functional (SCBIN & FCBIN), weighted structural and functional (SCWEI & FCWEI), binary structural weighted by function (SCFC), and binary functional weighted by structure (FCSC). Further details about constructing these connectomes can be found in the Methods, section 5.2.

Since the ten invariants are the building blocks of the graph space, the first question we asked was: to what extent are these invariants correlated? To answer this question, we computed the Spearman’s rank correlation coefficient (*ρ*) between every pair of invariants resulting in 10 *×* 10 correlation matrices, one for each model (see supplementary material, Figure S1). The same correlations were again computed at the RSN level. Figures 2a, 2b, and 2c illustrate the Spearman correlation patterns for each RSN of the following connectome models: weighted structural, weighted functional, and binary functional weighted by the structure. Results for the three other models can be seen in the Supplementary Material (Figures S2a, S2b, and S2c). These results chart an atlas of invariants where we can see which pairs of invariants carry redundant information, for the different RSNs and connectome models. The results show respectively positive and negative Spearman’s *ρ* values (respectively, in yellow and blue in Figure 2). As expected, for almost all models and RSNs the clustering coefficient and average shortest path length correlate negatively with all the other invariants and have strong positive correlation between them. The results at the RSN level show a general pattern of strong positive correlation between the clustering coefficient and average shortest path length; strong positive correlations between degree, betweenness centrality, closeness centrality, eigenvector centrality, within-module degree z-score, PageRank centrality, and subgraph; and moderately positive correlations between clustering coefficient and participation coefficient, for most of the RSNs. We also noted differences between RSNs within the same connectome models, for example the correlations of the betweenness centrality and the other invariants that change depending on the RSNs. These changes, which are more pronounced in the integrated SCFC and FCSC models, suggest that invariants of combined structural and functional connectomes yield less redundant information than the invariants of pure structural and functional networks.

**Figure 2:**
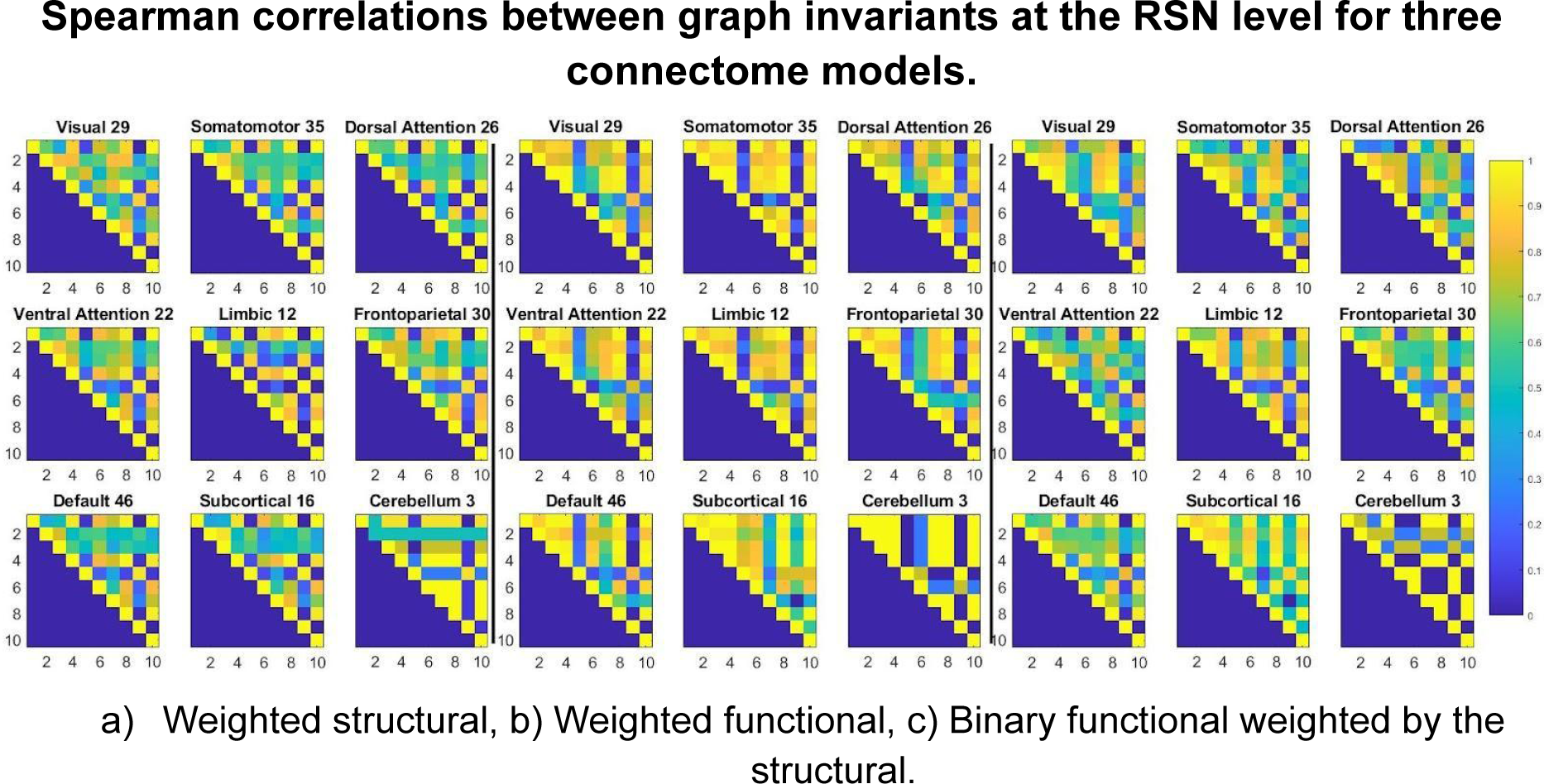
Spearman’s correlations computed average across subjects. Since the matrices are symmetric, only the upper triangular part is shown for visual simplicity. The nine matrices on the left, middle, and right represent the nine RSNs of the SCWEI, FCWEI, and FCSC, respectively. The number next to each RSN indicates the number of nodes from the whole network that belongs to the RSN. Invariant nomenclature; 1 = degree, 2 = betweenness centrality, 3 = closeness centrality, 4 = eigenvector centrality, 5 = clustering coefficient, 6 = participation coefficient, = 7 within-module degree z-score, 8 = PageRank centrality, 9 =average shortest path, and 10 = subgraph.

### Machine learning classification of lower order and higher order brain regions in multidimensional graph spaces

Next, we constructed multidimensional graph spaces for each model. Two examples are given in Figure 3, where we embed group-representative structural and functional connectomes into a 3D graph space built from degree centrality, participation coefficient, and within-module degree z-score invariants. From a visual inspection, brain regions tend to form distinct clusters in this space. For example, in figure 3b we can appreciate two distinct ‘bands’ of brain regions. The first band comprises areas from higher order brain networks, including the limbic, frontoparietal, default mode, subcortical, and cerebellum RSNs. The second band contains somatosensory areas belonging to the visual, somatomotor, dorsal, and ventral attention RSNs. We divided the RSNs into two categories for subsequent machine learning (ML) classification of brain regions into higher order (limbic, frontoparietal, default mode, subcortical and cerebellum RSNs) and lower order (visual, somatomotor, dorsal attention and central attention RSNs) regions. Qualitatively, higher order regions appear to have greater within-network connectivity and lower order regions have greater between-network connectivity (see Figure 3b).

**Figure 3:**
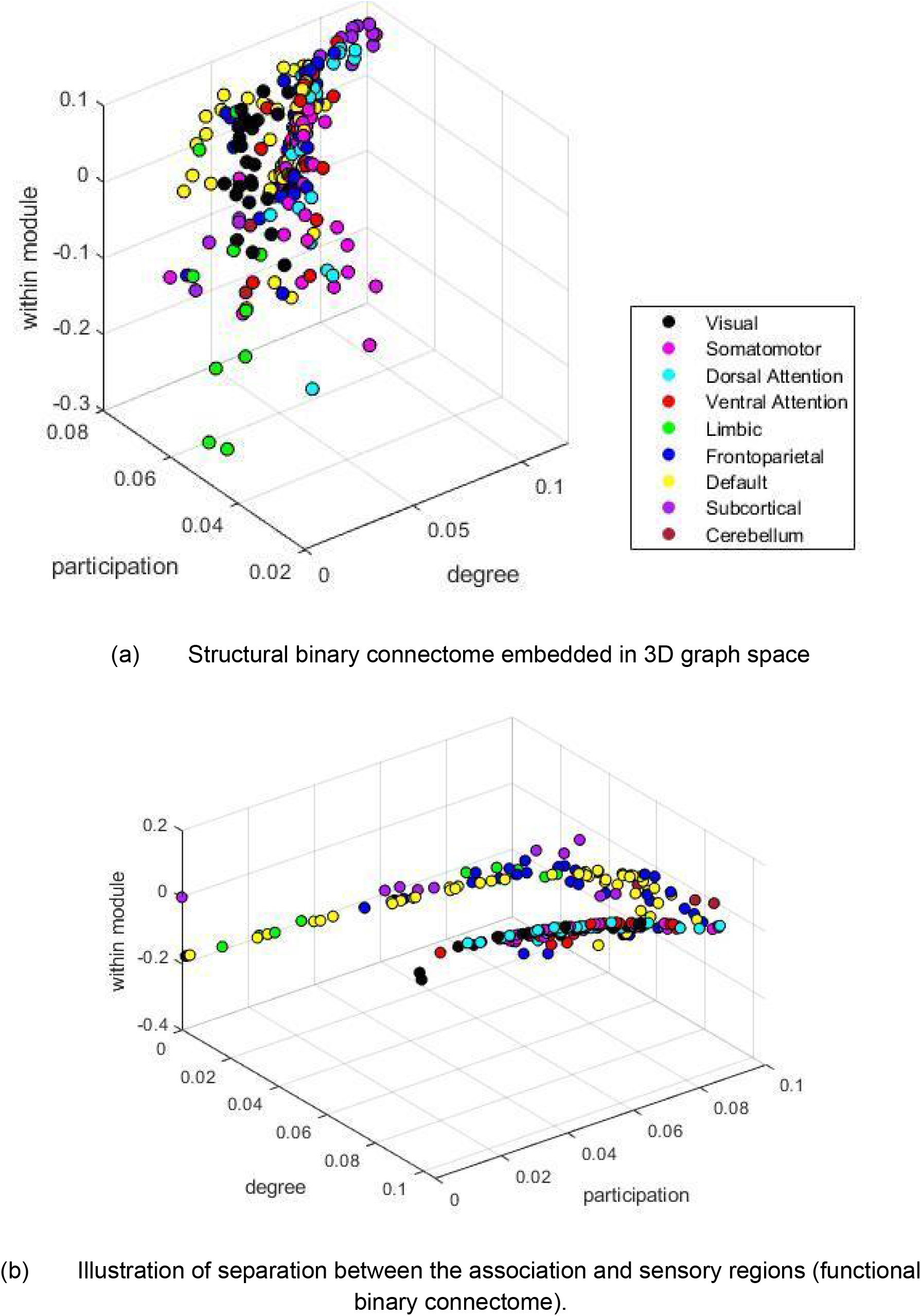
Examples of 3D graph spaces. Each dot represents a brain region colored according to the RSN it belongs to. The three axes of the 3D graph space correspond to the degree centrality, participation coefficient, and within-module degree z-score invariants.

We used supervised ML to test whether we could automatically classify brain regions into the two categories. We implemented the following classifiers: a binary support vector machine (SVM), a binary Gaussian kernel, and a binary linear classifier. The fitcsvm, fitckernel, and fitclinear Matlab functions [27–29] were used to create each classifier, respectively. For each model, all hyperparameters were optimized automatically by Matlab setting the OptimizeHyperparameters value to auto. In all cases, classifiers were trained with all the brain regions from 70 randomly selected subjects and then tested on all the brain regions of the remaining 30 subjects. Therefore, our trained models were tested on unseen data. In our case, training an algorithm on 70 randomly selected subjects means that all 70 subjects’ brain regions’ scores are used as input. In other words, each algorithm used a (70 ∗ 219 = 15330)*×n* matrix as input. The matrix’s rows correspond to the brain regions of the 70 randomly selected subjects, and the *n* columns represent the different invariant scores (*n* = 10 in a 10D graph space). The ML task is the following: given multiple brain regions’ graph-space coordinates (rows of the input matrix) and categories’ labels (higher order vs lower order regions), can the algorithm predict to which category the unseen brain areas belong? The unseen data used for testing is of shape (30 ∗ 219 = 6570)*×n*. Each algorithm was trained and tested on different graph space dimensions (2D, 3D, 10D) to see if the prediction increased with the dimensionality, based on the idea that a higher dimensional graph-space would yield more brain network information. An overview of our ML approach can be seen in Figure 4.

**Figure 4:**
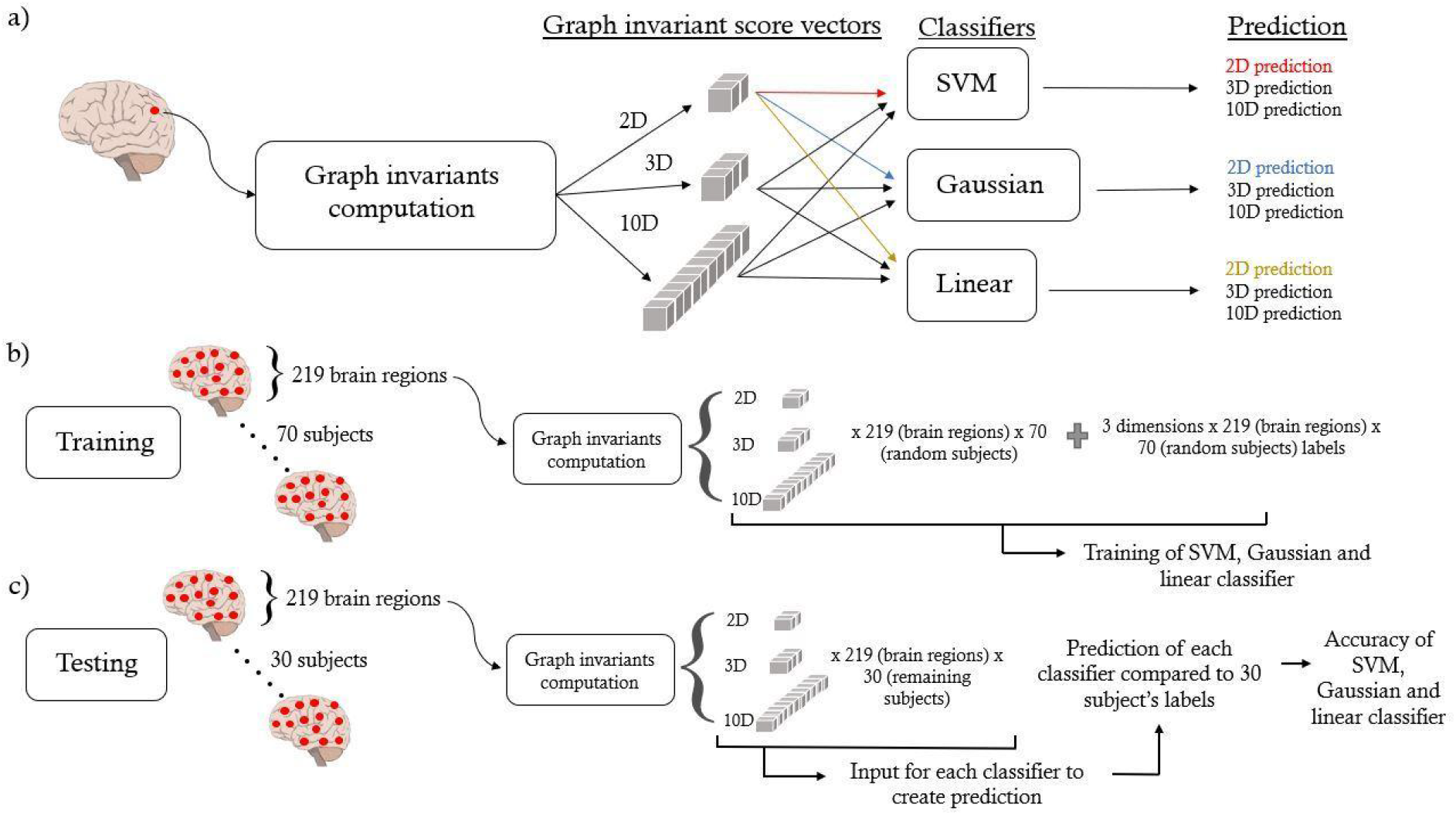
Schematic of machine learning analysis. (a) Overview of ML pipeline for one brain region. After computing all graph invariants, 2, 3, and all ten invariants of the brain region are used for each classifier resulting in a 2D, 3D, and 10D prediction. (b) Training pipeline. Each classifier was trained with 2, 3, and 10 graph invariants for all 219 brain regions of 70 randomly selected subjects. This resulted in 219 ∗ 70 = 15330 brain regions and labels for each dimension (2D, 3D, 10D). (c) Testing pipeline. 219 ∗ 30 = 6570 brain regions from the remaining 30 subjects were used as input for the classifiers to produce 6570 predictions. These predictions were compared to the labels to compute the classifier’s accuracy. This was done for all dimensions (2D, 3D, 10D).

We trained the models in 2D (degree and clustering), 3D (degree, clustering, and participation), and 10D spaces. The invariant’s choice for the 2D and 3D spaces was based on the correlation patterns (Supplementary Figure 1): two invariants with usually opposite correlations across all models (degree and clustering), and the same two plus an invariant showing lower correlations (participation), respectively. The accuracy of each classifier can be seen in Table 1 and was defined as the number of correctly classified brain regions divided by the total number of regions. Indirectly, these accuracy values indicate how suitable the graph space can be when trying to cluster brain areas. In our case, it directly measures how ML approaches can separate brain regions in abstract multidimensional graph spaces. As seen in Table 1, the highest test and train accuracy (87%) was achieved using a binary Gaussian classifier on the SCFC connectome model in a 10D metric space. The highest accuracies were achieved using all the ten invariants and ranged from 80% to 87%.

**Table 1:**
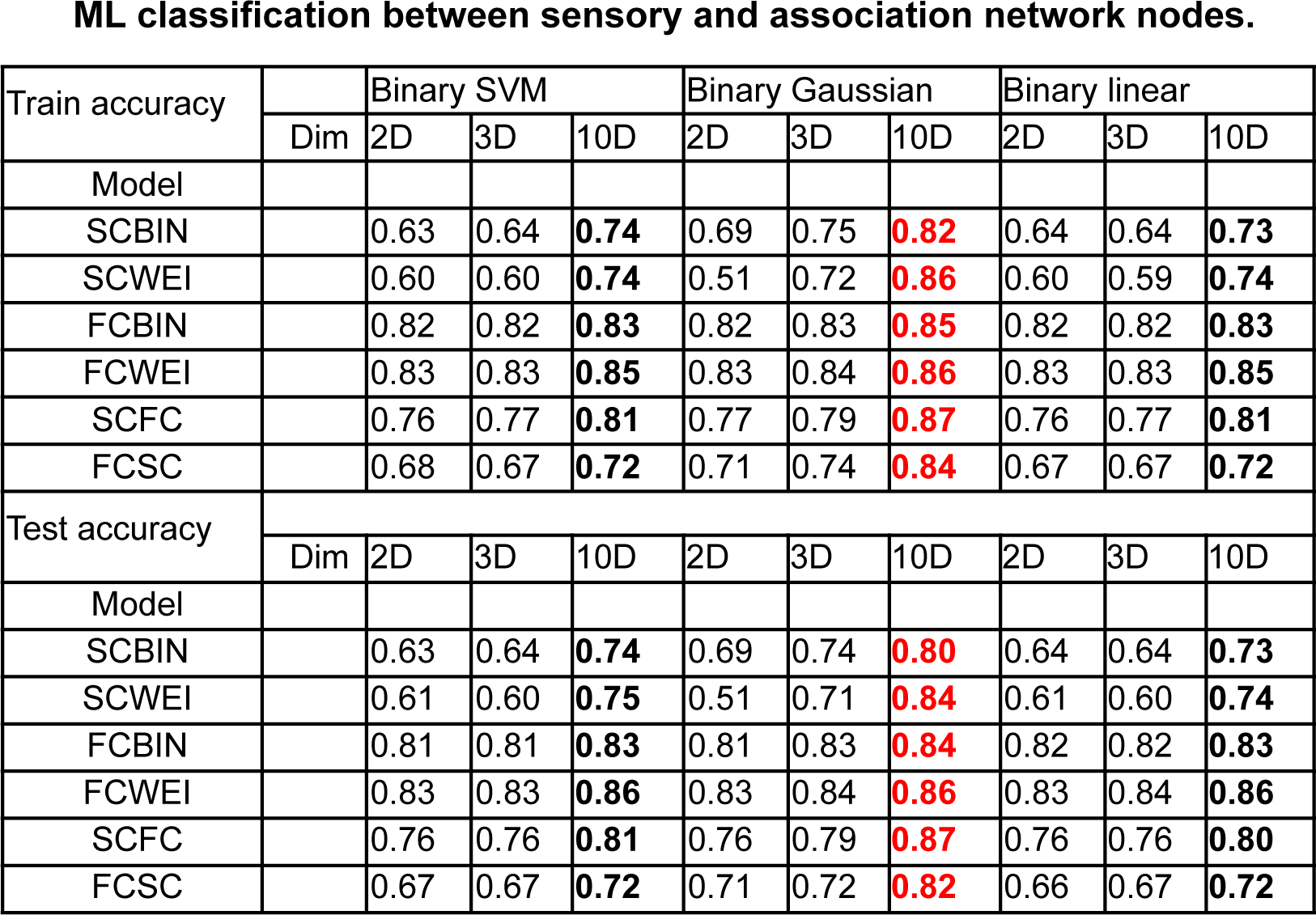
Bold numbers are the highest of the three dimensions for each algorithm and model. The red numbers represent the highest accuracies of all the corresponding rows.

| Train accuracy |  | Binary SVM |  |  | Binary Gaussian |  |  | Binary linear |  |  |
| --- | --- | --- | --- | --- | --- | --- | --- | --- | --- | --- |
|  | Dim | 2D | 3D | 10D | 2D | 3D | 10D | 2D | 3D | 10D |
| Model |  |  |  |  |  |  |  |  |  |  |
| SCBIN |  | 0.63 | 0.64 | **0.74** | 0.69 | 0.75 | **0.82** | 0.64 | 0.64 | **0.73** |
| SCWEI |  | 0.60 | 0.60 | **0.74** | 0.51 | 0.72 | **0.86** | 0.60 | 0.59 | **0.74** |
| FCBIN |  | 0.82 | 0.82 | **0.83** | 0.82 | 0.83 | **0.85** | 0.82 | 0.82 | **0.83** |
| FCWEI |  | 0.83 | 0.83 | **0.85** | 0.83 | 0.84 | **0.86** | 0.83 | 0.83 | **0.85** |
| SCFC |  | 0.76 | 0.77 | **0.81** | 0.77 | 0.79 | **0.87** | 0.76 | 0.77 | **0.81** |
| FCSC |  | 0.68 | 0.67 | **0.72** | 0.71 | 0.74 | **0.84** | 0.67 | 0.67 | **0.72** |
| Test accuracy |  |  |  |  |  |  |  |  |  |  |
|  | Dim | 2D | 3D | 10D | 2D | 3D | 10D | 2D | 3D | 10D |
| Model |  |  |  |  |  |  |  |  |  |  |
| SCBIN |  | 0.63 | 0.64 | **0.74** | 0.69 | 0.74 | **0.80** | 0.64 | 0.64 | **0.73** |
| SCWEI |  | 0.61 | 0.60 | **0.75** | 0.51 | 0.71 | **0.84** | 0.61 | 0.60 | **0.74** |
| FCBIN |  | 0.81 | 0.81 | **0.83** | 0.81 | 0.83 | **0.84** | 0.82 | 0.82 | **0.83** |
| FCWEI |  | 0.83 | 0.83 | **0.86** | 0.83 | 0.84 | **0.86** | 0.83 | 0.84 | **0.86** |
| SCFC |  | 0.76 | 0.76 | **0.81** | 0.76 | 0.79 | **0.87** | 0.76 | 0.76 | **0.80** |
| FCSC |  | 0.67 | 0.67 | **0.72** | 0.71 | 0.72 | **0.82** | 0.66 | 0.67 | **0.72** |

### Spatial comparison of brain regions of different subjects in a multi-dimensional graph space using Euclidean distance

The final analysis of this study was to investigate the distance between brain regions in the multidimensional graph spaces. Each brain region has a set of coordinates (its graph invariant scores), and we compute the distance between points using the Euclidean distance *d*:

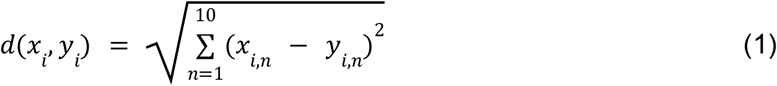

where *n* is the invariant; *x* and *y* two distinct subjects; *i* a specific brain region. Hence, *x*_*i*_ and *y*_*i*_ are vectors with ten values each, and *d(x*_*i*_, *y*_*i*_) represents the distance between the two brain regions, *x*_*i*_ and *y*_*i*_, embedded in the 10D graph space. Computing the Euclidean distance for each brain region between two different connectomes gives a vector of 219 values, each representing the Euclidean distance between the same brain region of different subjects. The distances between every pair of subjects were computed, allowing the comparison between different subjects. All these distances formed a 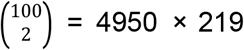 matrix for each connectome model. Averaging across all comparisons resulted in a total distance vector of 219 values for each model. These vectors represent the average inter-subject distance of each brain region, for each model. The results are shown in Figures 5a for the structural models, 5b for the functional models, and 5c for the mixed models. Darker colors indicate regions with larger inter-subject distance, thus representing the primary source of inter-subject variability. Furthermore, to mitigate the effect of potential outliers, we computed for each brain region the ratio between the mean and the standard deviation of its pair-wise inter-subject distances. These results can be seen in the supplementary material (Supplementary Figure 3).

**Figure 5:**
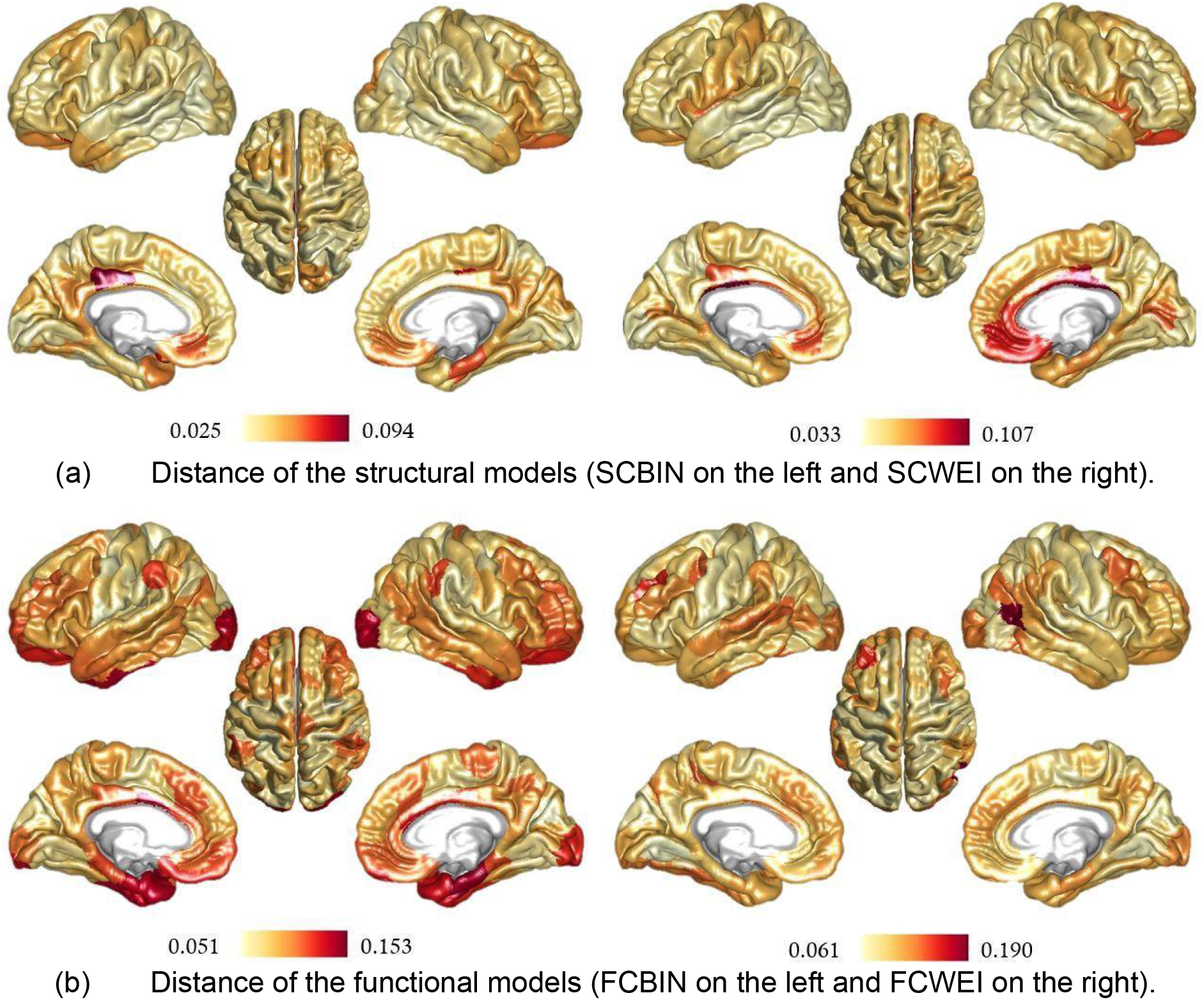
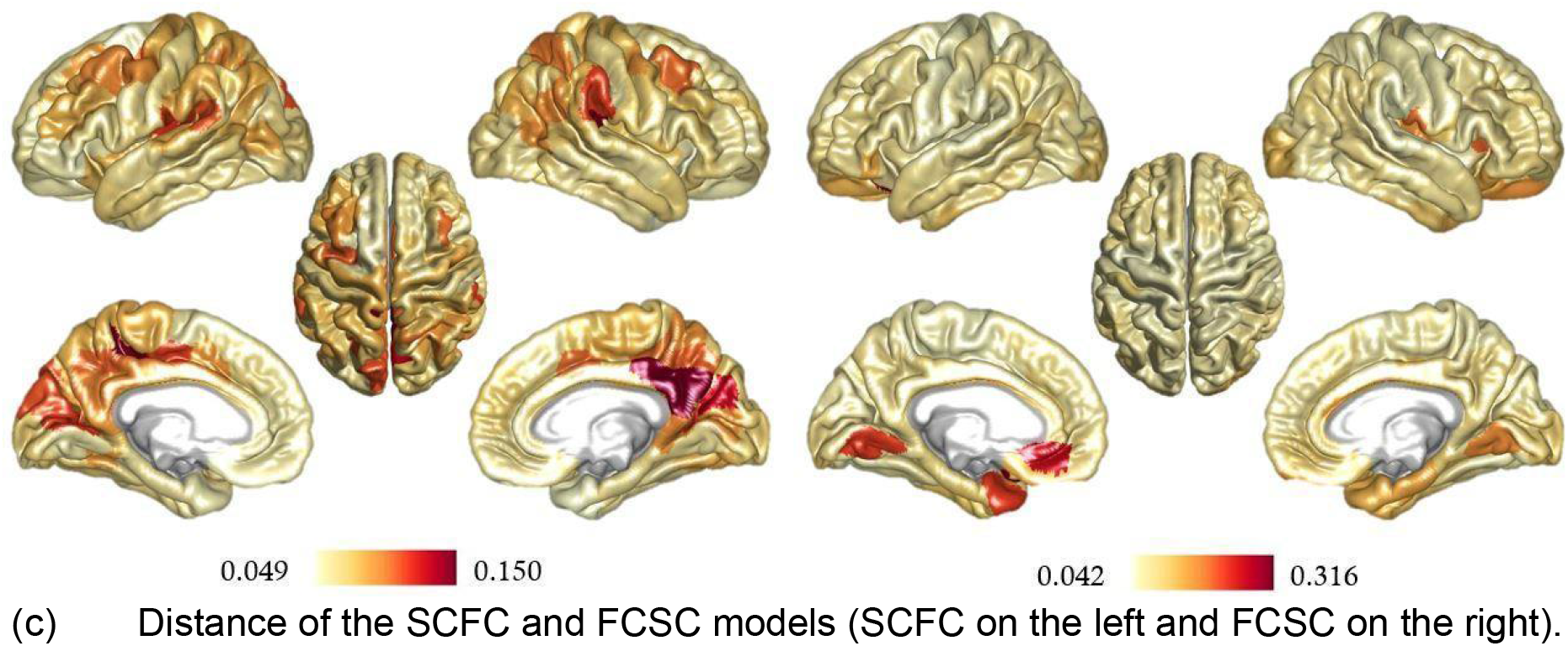
Inter-subject nodal distance in 10-dimensional graph spaces[AG1]. Brain regions exhibiting a high inter-subject distance are in red, and brain regions showing a low inter-subject distance are in white. These brain maps represent the inter-subject variability from a graph space perspective, with red regions being highly subject-specific and white regions having similar invariant scores across subjects.

For the structural models, regions of the default mode network exhibit the highest inter-subject distances in the binary connectomes, and temporal regions in the weighted connectomes. The pattern is less clear in the functional models; nevertheless, the frontoparietal and temporal areas always yield relatively high inter-subject distances, suggesting that these regions carry the largest proportion of inter-individual variability in nodal topological embedding within multidimensional graph spaces. The highest inter-subject distances concern the functional and mixed models (FCBIN, FCWEI, SCFC, and FCSC) while the lowest distances concern the structural models (SCBIN and SCWEI). This result suggests that the functional information and the structure-function mixed information tend to better differentiate subjects than the structural information alone.

After investigating inter-individual distances between homologous brain regions, we computed the average inter-individual distance of all 219 brain regions to get a single global distance measure between connectome pairs. This resulted in 4950 global distances, one between every pair of subjects, which were then averaged. This value was computed at multiple network densities to determine how fast these distances would increase or decrease as a function of density for each model. The results for each independent model and all models together can be seen in the supplementary material (Figures 4, and 5). Here, the density of our connectomes is just the amount of strongest connections kept concerning all connections. These results are to be taken as proof that a single global distance between different connectomes can be computed. The implication of these global distances and curves, in our results, will not be discussed here since, some graph invariants can return aberrant values at low densities. More work will be needed to use this global connectome distance better.

## Discussion

In this work, we proposed a new graph space to study connectomes. Our method could be conceptualized as a generalization of previous 2D and 3D graph invariant approaches [22, 23]. Our results provide the following new information; 1) graph invariants are correlated or anticorrelated at different levels (i.e., there exists specific information between invariants); 2) machine learning algorithms can separate information in multidimensional graph spaces, especially in high dimensional spaces; 3) embedding brain regions in a Euclidean graph space provides a mathematical definition of distance between brain regions at multiple scales; and 4) multidimensional graph spaces offer a new way to compare two, or more, connectomes from different subjects. We believe this new methodology of graph spaces could be helpful in computational and clinical neurosciences.

The invariants’ correlation patterns showed differences in graph invariants at the brain subnetwork level and for different brain network models. These surprising patterns imply that each RSN should be studied with special care regarding the choice of invariants. Each RSN is known to be involved in different cognitive and biological functions [30–32]. However, the Spearman results illustrate this fact from the existing networks. These changes in patterns make sense regarding the differences between connectome models (SCBIN, SCWEI, FCBIN, etc.), as illustrated in Supplementary Figure 1. Either the structure of the networks is different (the binary structural and functional networks of the brain are known to be different [7, 8]), or the different types of weights (physical vs. topological distances) are different. Both can influence the computation of the invariants. The more particular fact is the contrast in correlation patterns between RSNs (Figure 2). As previously mentioned, these networks are known to be distinct. The divergent correlation patterns between RSNs underline this fact via quantitative nodal graph importance measurements. This suggests that a brain region’s importance must be measured considering its sub-network environment. In other words, the relevance of a brain region in its RSN should not always be compared with areas from other RSNs using the same invariants. Results from Figures 2a, 2b, and 2c can be considered maps for navigators trying to dismantle the RSN roads sailing with invariants as their compass.

The graph space has proven to be well-suited for ML applications. In our work, we were able to achieve 87% of accuracy when classifying brain regions between somatosensory vs. associate network areas. Our methodology offers a new approach to studying brain areas because they can naturally form clusters in our space. This brings a new unique network classification of these areas instead of a biological one. It supports the idea that the graph space allows network nodes to isolate, cluster, or spread themselves spontaneously without any prior hypothesis. As all testing accuracies were relatively high, our results also reinforce the idea that the graph space could be used with clinical data to build early and noninvasive diagnostics of neurological diseases. All three different ML regimes tested achieved decent results (Table 1), and almost all the best results came from the Gaussian classifier. This indicates that nonlinear-shaped algorithms are better suited for brain network classification in graph spaces. Another aspect worth remarking is that our ML approach was computed on whole brain connectomes but can also be used with subnetworks. For example, if a disease is known to target specific brain areas, only networks involving these areas can be studied via the graph space. Thus, this space can be used with networks representing a different level of brain complexity (whole brain, subnetworks, fewer or more nodes for the same scale, etc.), showing the adaptability and flexibility of the proposed tool to match researchers’ needs. All our accuracies also support the idea of increasing information with the degree of dimension. This shifts the way brain networks are studied since, as previously mentioned, most work regarding connectomes is done in 2 or 3D.

Our current ML approach has some limitations, the biggest being that K-fold cross-validation should be implemented to reach better accuracy since our results come from hold-out cross-validation. The number of subjects used (100) could also be increased, and more sophisticated ML methods could be used, such as Neural Networks for example. Nevertheless, we show preliminary evidence that more extensive information, understanding, and better classification could come from these networks when more dimensions are in place.

Finally, we proposed a new distance measure in the graph space to compare single brain regions, Figure 5 and Supplementary Figure 3, and the whole brain comparison, Supplementary Figure 4 and 5. Connectomes of different subjects express differences between them [33, 34]. However, the graph space offers us a new angle to explore discrepancies between connectomes, regarding the importance of each brain region, via multiple network algorithms. Therefore, the distance across subjects’ results expresses how far brain regions are from a network perspective. More specifically, if the distance between the same pair of regions from different subjects is high, this region doesn’t yield the same importance for both subjects. From our results (Figure 5 and Supplementary Figure 3), both structural and functional distances should be considered to understand the brain since their distance pattern is different (they represent different information). The structural distances yielded high values in the default network areas and low values in frontoparietal and visual networks. Since the default network is widely spread across the brain [30, 35, 36], it makes sense that it could differ structurally (in length or strength) between different people. The functional distances appeared to be highest in the frontoparietal and temporal areas and lowest in the somatomotor and dorsal networks. The frontoparietal regions are known to be involved in complex cognitive activities [37]; thus, it is reasonable that these regions are subject-specific and express high distances from each other in fMRI data. Whole distance results are not very easy to interpret since we lose some information having only one distance value.

Nevertheless, a global trend is that all networks tend to be closer to each other at high network density and the opposite at low density. This makes sense since, as mentioned previously, some invariants return aberrant values at low densities. Also, functional networks exhibit higher distances compared to structural models. This expresses that functional brain information is more subject-specific than structural data, which was expected since every human has the same global structure of the brain. In other words, the actual physical distance used in the structural connectomes is more similar across subjects than the topological functional distance, which is expected from a biological point of view.

This (single region and whole brain) distance measure offers a new tool to identify how brain regions are spatially close. If a template of a healthy brain could be made, this distance could be used to see how far certain brain regions, known to be involved in neurological diseases, are from the template. Reversing the question, if a model of a brain with a specific disease can be created, we could measure how close the brain regions of patients are to it in a harmless and non-invasive way. This distance measure could then yield brain maps similar to the ones from Figure 5 and Supplementary Figure 3 for each neurological disease. Following this, a single brain region distance map could be generated and compared between young vs. elder subjects to assess longitudinal brain changes.

Future work will be needed to exploit all the graph space possibilities. Regarding the ML approaches, analyses of clinical data would help identify cluster-specific brain regions involved in neurological diseases non-incisively. Neural networks could also be investigated to see if they are more efficient than our method of identifying brain data patterns embedded in high dimensional graph space. Also, ML analyses of subnetworks can offer a new perspective on identifying network differences and patterns extractable from and across them. From a biological point of view, connectomes of different species can be embedded and compared in a graph space. Investigating how intelligent or primal species, e.g., big vs. small animal brain networks, spread in the graph space could lead to a spatial gradient of species. Another relevant question for future work is which invariants are redundant in gaining new information. We argue here that high-dimensional spaces are well suited for studying connectomes, but does each new dimension (each invariant) give us more information than we already had in lower-dimensional spaces? Two types of answers can be considered; 1) Some invariants do not improve the knowledge of the connectome in a graph space sense, leading to a debate over which invariants are the best. Alternatively, 2) any new centrality computed can either increase the amount of global information or equal it but never decrease it. We will not tackle this question here, since our work aims to be a proof of concept regarding the graph space rather than an explanation of all the technical details, but this aspect should be analyzed in future work.

As a final word, we explore here the graph space in the context of neuroscience. However, as mentioned previously, it can be used with any network; therefore, it is not just a fruitful tool in computational neuroscience but could lead to novel findings in any other field involving the use of networks.

## Methods

### Data set and preprocessing

This study used a Human Connectome Project dataset of structural and functional MRI data from 100 unrelated subjects [26]. All diffusion data were processed following the main MRtrix guidelines [41] with two modifications; 1) the track seeding was set to 20 million tracts, and 2) the algorithm ”spherical deconvolution informed filtering of tractograms” 2.0 [43] was used.

The fMRI data included four sessions of resting-state acquired for each subject (two per day with opposed phase encoding direction), each composed of time series of 1200 acquired volumes (TR=0.72s between volumes) that were preprocessed as in [33]. Briefly, preprocessing steps included linear and quadratic detrending, removal of motion regressors and their first derivatives, removal of white matter, cerebrospinal fluid average signals and their first derivatives, and a bandpass filter in the range of 0.01-0.15 Hz.

### Connectomes

The 200 cortical regions of the Schaefer parcellation [39], plus 19 subcortical and cerebellum regions (as in [44]) were used to create the connectomes.

The edges of both structural connectomes (SCBIN & SCWEI) represent anatomical connections between the brain regions. Two graphs were created using two metrics as edges: fiber length and the number of streamlines [40]. The fiber length matrices represent the distance separating two brain regions in millimeters, and the number of streamline matrices expresses the strength of the connection between different brain areas. A consensus mask across all subjects was computed by binarizing the number of streamline matrices. This resulted in a matrix where the *ij*^*th*^ element is one if there is a connection between brain regions *i* and *j* across all subjects; otherwise, it is zero. Next, the average of all 100 fiber length matrices was multiplied with the consensus mask, giving an average masked fiber length matrix. This matrix is a structural connectome created using all subjects’ structural information. The density of this average structural connectome was used to obtain single-subject structural connectomes, where individual fiber length matrices were thresholded to keep only more robust connections and match the average connectome density. The SCWEI model considers these single-subject fiber length matrices, while the SCBIN model considers the same matrices but binarized.

For the functional connectomes (FCBIN & FCWEI), the edges do not represent a physical connection but a statistical correlation between fMRI signals of different brain areas. The pairwise correlations between regional average time series were computed and considered as each individual’s edges of a functional connectome. The information these matrices convey does not refer to physical links between nodes but rather how similar their activity pattern is. Conversely, if the activity of two brain regions is desynchronized, their correlation will be low. The absolute value of each element was computed, and the density of the average masked fiber length matrix was again used as a threshold to keep only the most robust connections. Furthermore, the FCWEI model is the single-subject matrix with the original correlation numbers, while the FCBIN model is the same matrix but thresholded to have a binary matrix.

The SCFC model was created by multiplying the binary structural matrix by the weighted functional matrix for each subject. This model represents a structural graph with weights coming from the functional data, so the correlation of their activity pattern weighs the anatomical connections between brain regions. The FCSC model is the opposite, i.e., constructed by multiplying each subject’s binary functional matrix with the average fiber length matrix, giving us a functional graph with weights from the anatomical data. In this case, the correlation between brain regions is weighted by the physical distance separating them.

### Graph invariants

Let *G* be a graph with *n* vertices, |*V*| = *n*. The adjacency matrix *A* of *G* is an *n x n* symmetric matrix where each element *a* is defined as,

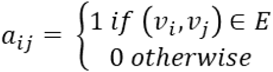

A graph invariant is an algorithm that takes as input a graph, *G*, and computes a score, *s*_*i*,_ for each vertex belonging to *V*. Each vertex score represents how vital this vertex is in the whole graph. The ten following invariants were used in this study:

### Degree

The degree invariant, *C*_*D*_, of a vertex *i* is defined as the number of neighbors of *i. C*_*D*_*(i)* = *deg(v*_*i*_).

### Betweenness

The betweenness invariant, *C*_*b*_, of a vertex *i* is the ratio of the shortest paths connecting any pair of vertices, *s* and *t*, that passes through *i* divided by the total number of shortest paths between *s* and *t*. 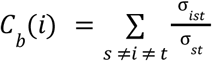 where *σ*_*sit*_ is the number of shortest paths from *s* to *t* that passes through *i*, and *σ*_*st*_ is the total number of shortest paths from *s* to *t*.

### Closeness

The closeness,*C*_*c*_, of a vertex is the inverse sum of the length of the shortest paths connecting the vertex to all other vertices in the graph.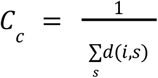 where *d(i, s)* is the distance between vertices

### Eigenvector

The eigenvector invariant, *C*_*e*_, is based on the idea that if a vertex *v* is linked to many vertices with a high eigenvector score, it will also have a high eigenvector score, and vice versa if it is linked with lower scores.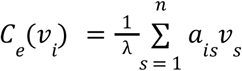, where *λ* is the highest eigenvalue of the adjacency matrix *A*.

### Clustering

The clustering coefficient, *C*_*clu*_, of a vertex *i*, represents how much the vertices connected to *i* are linked together. 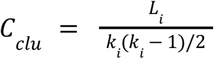, where *L*_*i*_ is the number of edges between the neighbors of *i* and *k*_*i*_ is the degree of *i*.

### Participation

The participation coefficient, *C*_*p*_, reflects how much a vertex *i* is linked with vertices of its module or vertices from other modules. Modules can be thought of as clusters of vertices that are highly interlinked. 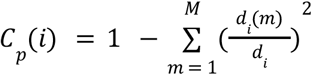, where *M* is the set of all the modules, *d*_*i*_ is the degree of vertex *i*, and *d*_*i*_*(m)* is the degree between vertex *i*, and all vertices in module *m*.

### Within-module degree z-score

The within-module degree z-score, *C*_*z*_, of a vertex *i*, is the z-score of *i* within its module. It is related to the degree of a vertex but only within its module.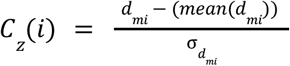, where *d* is the degree of vertex 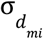 in its module *m*_*i*_, *mean(d*_*mi*_) is the mean degree of *i* within it’s module *m*_*i*_, and 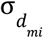 is the standard deviation of the degree of *i* within its’ module.

### PageRank

The PageRank Google’s invariant, *C*_*pr*_, is a variation of the eigenvector invariant but follows the same idea of self-reference. The PageRank score of a vertex *i* can be thought of as the amount of time a random walker would spend on *i* concerning the whole graph and with a damping factor specifying the amount of time the random walker will step on a neighborhood vertex *i*.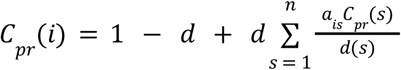, where *d* is the damping factor, usually 0.85, and *d(s)* is the degree of vertex *s*.

### Average shortest path

The average shortest path invariant, *C*_*asp*_, is a measure of the efficiency of a vertex in transferring information in the whole network. It can be defined as the reciprocal of nodal efficiency. 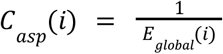, where 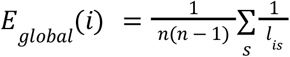 is the global efficiency of vertex *i* with *l*_*is*_ being the shortest path between vertices *i* and *s*.

### Subgraph

The subgraph invariant, *C*_*s*_, of a vertex *i* is a weighted sum of a closed walk that can be of different lengths, starting and ending at *i*. 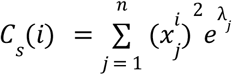, where *x*_*j*_ is the *j*th eigenvector of *A, x*^*i*^ is the *i*th component of *x*_*j*_, and *λ*_*j*_ is the *j*th eigenvector.

All the analyses and invariants were computed on Matlab using the Brain Connectivity Toolbox [42], and after computation, all invariants were individually normalized.

## Acknowledgments

MGP was supported by the CIBM Center for Biomedical Imaging, a Swiss research center of excellence founded and supported by Lausanne University Hospital (CHUV), University of Lausanne (UNIL), Ecole polytechnique fédérale de Lausanne (EPFL), University of Geneva (UNIGE) and Geneva University Hospitals (HUG).

## References

1. Kleinberg, J. & Lawrence, S. The Structure of the Web. Science 294. Publisher: American Association for the Advancement of Science, 1849–1850. https://www.science.org/doi/full/10.1126/science.1067014 (2022) (Nov. 30, 2001).

2. Jordán, F., Benedek, Z. & Podani, J. Quantifying positional importance in food webs: A comparison of centrality indices. Ecological Modelling 205, 270–275. issn: 0304-3800. https://www.sciencedirect.com/science/article/pii/S0304380007001184 (2022) (July 10, 2007).

3. Park, H. W. Hyperlink network analysis: A new method for the study of social structure on the web. Connections 25, 49–61 (2003).

4. Pedersen, M., Omidvarnia, A., Shine, J. M., Jackson, G. D. & Zalesky, A. Reducing the influence of intramodular connectivity in participation coefficient. Network Neuroscience 4, 416–431. issn: 2472-1751. https://doi.org/10.1162/netn_a_00127 (2022) (Apr. 1, 2020).

5. Das, K., Samanta, S. & Pal, M. Study on centrality measures in social networks: a survey. Social Network Analysis and Mining 8, 13. issn: 1869-5469. https://doi.org/10.1007/s13278-018-0493-2 (2022) (Feb. 28, 2018).

6. Du, Y. et al. A new closeness centrality measure via effective distance in complex networks. Chaos: An Interdisciplinary Journal of Nonlinear Science 25. Publisher: American Institute of Physics, 033112. issn: 1054-1500. https://aip.scitation.org/doi/full/10.1063/1.4916215 (2022) (Mar. 2015).

7. Fornito, A., Zalesky, A. & Bullmore, E. Fundamentals of Brain Network Analysis 496 pp. isbn: 978-0-12-408118-5 (Academic Press, Mar. 4, 2016).

8. Sporns, O. Networks of the Brain Google-Books-ID: 9LlNEAAAQBAJ. 433 pp. isbn: 978-0-262-52898-6 (MIT Press, Feb. 12, 2016).

9. Bullmore, E. & Sporns, O. Complex brain networks: graph theoretical analysis of structural and functional systems. Nature Reviews Neuroscience 10. Number: 3 Publisher: Nature Publishing Group, 186–198. issn: 1471-0048. https://www.nature.com/articles/nrn2575 (2022) (Mar.2009).

10. Simas, T., Chavez, M., Rodriguez, P. & Diaz-Guilera, A. An algebraic topological method for multimodal brain networks comparisons. Frontiers in Psychology 6. issn: 1664-1078. https://www.frontiersin.org/article/10.3389/fpsyg.2015.00904 (2022) (2015).

11. Farahani, F. V., Karwowski, W. & Lighthall, N. R. Application of Graph Theory for Identifying Connectivity Patterns in Human Brain Networks: A Systematic Review. Frontiers in Neuroscience 13. issn: 1662-453X. https://www.frontiersin.org/article/10.3389/fnins.2019.00585 (2022) (2019).

12. Wang, J., Zuo, X. & He, Y. Graph-based network analysis of resting-state functional MRI. Frontiers in Systems Neuroscience 4. issn: 1662-5137. https://www.frontiersin.org/article/10.3389/fnsys.2010.00016 (2022) (2010).

13. Stam, C. J. Modern network science of neurological disorders. Nature Reviews Neuroscience 15. Number: 10 Publisher: Nature Publishing Group, 683–695. issn: 1471-0048. https://www.nature.com/articles/nrn3801 (2022) (Oct. 2014).

14. Avena-Koenigsberger, A., Goni, J., Sole, R. & Sporns, O. Network morphospace. Journal of The Royal Society Interface 12. Publisher: Royal Society, 20140881. https://royalsocietypublishing.org/doi/full/10.1098/rsif.2014.0881 (2022) (Feb. 6, 2015).

15. Goñi, J. et al. Exploring the Morphospace of Communication Efficiency in Complex Networks. PLOS ONE 8. Publisher: Public Library of Science, e58070. issn: 1932-6203. https://journals.plos.org/plosone/article?id=10.1371/journal.pone.0058070 (2022) (Mar. 7, 2013).

16. Zhao, S., Rangaprakash, D., Liang, P. & Deshpande, G. Deterioration from healthy to mild cognitive impairment and Alzheimer’s disease mirrored in corresponding loss of centrality in directed brain networks. Brain Informatics 6, 8. issn: 2198-4026. https://doi.org/10.1186/s40708-019-0101-x (2022) (Dec. 2, 2019).

17. Bassett, D. S. & Bullmore, E. T. Human Brain Networks in Health and Disease. Current opinion in neurology 22, 340–347. issn: 1350-7540. https://www.ncbi.nlm.nih.gov/pmc/articles/PMC2902726/ (2022) (Aug. 2009).

18. Thomas Yeo, B. T. et al. The organization of the human cerebral cortex estimated by intrinsic functional connectivity. Journal of Neurophysiology 106. Publisher: American Physiological Society, 1125–1165. issn: 0022-3077. https://journals.physiology.org/doi/full/10.1152/jn.00338.2011 (2022) (Sept. 2011).

19. Sporns, O. Graph theory methods: applications in brain networks. Dialogues in Clinical Neuro-science 20, 111–121. issn: 1958-5969 (June 2018).

20. Mantzaris, A. V. et al. Dynamic network centrality summarizes learning in the human brain. Journal of Complex Networks 1, 83–92. issn: 2051-1310. https://doi.org/10.1093/comnet/cnt001 (2022) (June 1, 2013).

21. Kuhnert, M.-T., Geier, C., Elger, C. E. & Lehnertz, K. Identifying important nodes in weighted functional brain networks: A comparison of different centrality approaches. Chaos: An Interdisciplinary Journal of Nonlinear Science 22. Publisher: American Institute of Physics, 023142. issn: 1054-1500. https://aip.scitation.org/doi/10.1063/1.4729185 (2022) (June 2012).

22. Joyce, K. E., Laurienti, P. J., Burdette, J. H. & Hayasaka, S. A New Measure of Centrality for Brain Networks. PLOS ONE 5. Publisher: Public Library of Science, e12200. issn: 1932-6203. https://journals.plos.org/plosone/article?id=10.1371/journal.pone.0012200 (2022) (Aug. 16, 2010).

23. Zuo, X.-N. et al. Network Centrality in the Human Functional Connectome. Cerebral Cortex 22, 1862–1875. issn: 1047-3211. https://doi.org/10.1093/cercor/bhr269 (2022) (Aug. 1, 2012).

24. Van den Heuvel, M. P. & Sporns, O. A cross-disorder connectome landscape of brain dysconnectivity. Nature Reviews Neuroscience 20. Number: 7 Publisher: Nature Publishing Group, 435–446. issn: 1471-0048. https://www.nature.com/articles/s41583-019-0177-6 (2022) (July 2019).

25. Sun, D., Haswell, C. C., Morey, R. A. & Bellis, M. D. D. Brain structural covariance network centrality in maltreated youth with PTSD and in maltreated youth resilient to PTSD. Development and Psychopathology 31. Publisher: Cambridge University Press, 557–571. issn: 0954-5794, 1469-2198. https://www.cambridge.org/core/journals/development-and-psychopathology/article/abs/brain-structural-covariance-network-centrality-in-maltreated-youth-with-ptsd-and-in-maltreated-youth-resilient-to-ptsd/8A9C2DC433527D518FA450A34FC3B25A (2022) (May 2019).

26. Human Connectome Project — Mapping the human brain connectivity http://www.humanconnectomeproject.org/ (2022).

27. Train support vector machine (SVM) classifier for one-class and binary classification - MATLAB fitcsvm - MathWorks Switzerland https://ch.mathworks.com/help/stats/fitcsvm.html (2022).

28. Fit binary Gaussian kernel classifier using random feature expansion - MATLAB fitckernel - Math- Works Switzerland https://ch.mathworks.com/help/stats/fitckernel.html (2022).

29. Fit binary linear classifier to high-dimensional data - MATLAB fitclinear - MathWorks Switzerland https://ch.mathworks.com/help/stats/fitclinear.html (2022).

30. Seitzman, B. A., Snyder, A. Z., Leuthardt, E. C. & Shimony, J. S. The State of Resting State Networks. Topics in Magnetic Resonance Imaging 28, 189–196. issn: 1536-1004. https://journals.lww.com/topicsinmri/Fulltext/2019/08000/The_State_of_Resting_State_Networks.1.aspx (2022) (Aug. 2019).

31. Damoiseaux, J. S. et al. Consistent resting-state networks across healthy subjects. Proceedings of the National Academy of Sciences 103. Publisher: Proceedings of the National Academy of Sciences, 13848–13853. https://www.pnas.org/doi/full/10.1073/pnas.0601417103 (2022) (Sept. 12, 2006).

32. Heuvel, M. P. v. d. & Sporns, O. An Anatomical Substrate for Integration among Functional Networks in Human Cortex. Journal of Neuroscience 33. Publisher: Society for Neuroscience Section: Articles, 14489–14500. issn: 0270-6474, 1529-2401. https://www.jneurosci.org/content/33/36/14489 (2022) (Sept. 4, 2013).

33. Van De Ville, D., Farouj, Y., Preti, M. G., Liégeois, R. & Amico, E. When makes you unique: Temporality of the human brain fingerprint. Science Advances 7. Publisher: American Association for the Advancement of Science, eabj0751. https://www.science.org/doi/10.1126/sciadv.abj0751 (2022) (Oct. 15, 2021).

34. Varoquaux, G. & Craddock, R. C. Learning and comparing functional connectomes across subjects. NeuroImage. Mapping the Connectome 80, 405–415. issn: 1053-8119. https://www.sciencedirect.com/science/article/pii/S1053811913003340 (2022) (Oct. 15, 2013).

35. Van den Heuvel, M. P. & Hulshoff Pol, H. E. Exploring the brain network: A review on resting-state fMRI functional connectivity. European Neuropsychopharmacology 20, 519–534. issn: 0924-977X. https://www.sciencedirect.com/science/article/pii/S0924977X10000684 (2022) (Aug. 1, 2010).

36. Shulman, G. L. et al. Common Blood Flow Changes across Visual Tasks: II. Decreases in Cerebral Cortex. Journal of Cognitive Neuroscience 9, 648–663. issn: 0898-929X. https://doi.org/10.1162/jocn.1997.9.5.648 (2022) (Oct. 1, 1997).

37. Marek, S. & Dosenbach, N. U. F. The frontoparietal network: function, electrophysiology, and importance of individual precision mapping. Dialogues in Clinical Neuroscience 20, 133–140. issn: 1294-8322. https://www.ncbi.nlm.nih.gov/pmc/articles/PMC6136121/ (2022) (June 2018).

38. Lab :: MIP https://miplab.epfl.ch/ (2022).

39. CBIG/stable projects/brain parcellation/Schaefer2018 LocalGlobal at master · ThomasYeoLab/CBIG GitHub. https://github.com/ThomasYeoLab/CBIG (2022).

40. Griffa, A. et al. The evolution of information transmission in mammalian brain networks Pages: 2022.05.09.491115 Section: New Results. May 10, 2022. https://www.biorxiv.org/content/10.1101/2022.05.09.491115v1 (2022).

41. Welcome to the MRtrix3 user documentation! — MRtrix 3.0 documentation https://mrtrix.readthedocs.io/en/latest/ (2022).

42. Brain Connectivity Toolbox https://sites.google.com/site/bctnet/ (2022).

43. Smith, R. E., Tournier, J.-D., Calamante, F. & Connelly, A. SIFT2: Enabling dense quantitative assessment of brain white matter connectivity using streamlines tractography. NeuroImage 119, 338–351. issn: 1053-8119. https://www.sciencedirect.com/science/article/pii/S1053811915005972 (2023) (Oct. 1, 2015).

44. Griffa, A., Amico, E., Liégeois, R., Van De Ville, D. & Preti, M. G. Brain structure-function coupling provides signatures for task decoding and individual fingerprinting. NeuroImage 250, 118970. issn: 1053-8119. https://www.sciencedirect.com/science/article/pii/S1053811922000994 (2023) (Apr. 15, 2022).

