## Supplementary material for "Connectome embedding in multidimensional graph-invariant spaces"

November 2022

[1] Neuro-X Institute, Ecole Polytechnique Fédérale De Lausanne (EPFL), Geneva, Switzerland

[2] Department of Radiology and Medical Informatics, University of Geneva (UNIGE), Geneva, Switzerland

[3] CIBM Center for Biomedical Imaging, Switzerland

[4] Leenaards Memory Center, Lausanne University Hospital and University of Lausanne, Lausanne, Switzerland

[5] Department of Psychology and Neuroscience, Auckland University of Technology, New Zealand

**Spearman correlations between graph invariants at the whole connectome level.**

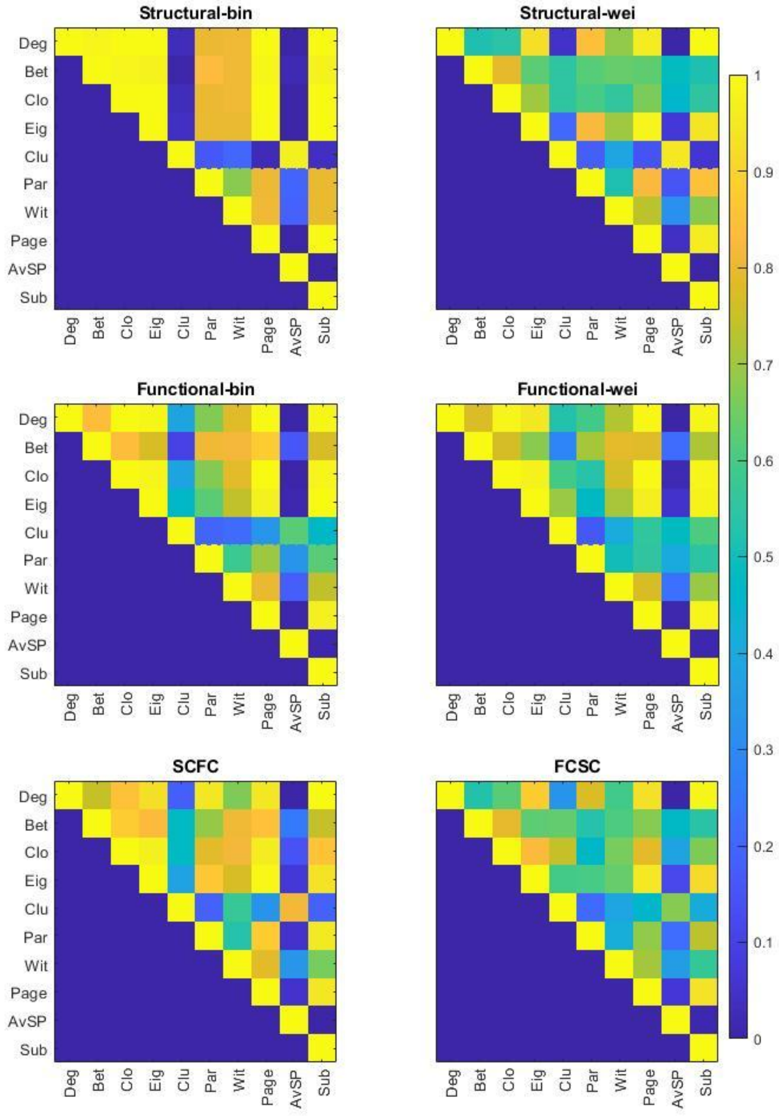


**Supplementary Figure 1:** Spearman's correlations at whole connectome level, for each model. All correlations were computed on the average level. Since the matrices are symmetric, only the upper triangular part is shown for visual simplicity.

**Spearman correlations between graph invariants at RSN level for three different connectome models.**

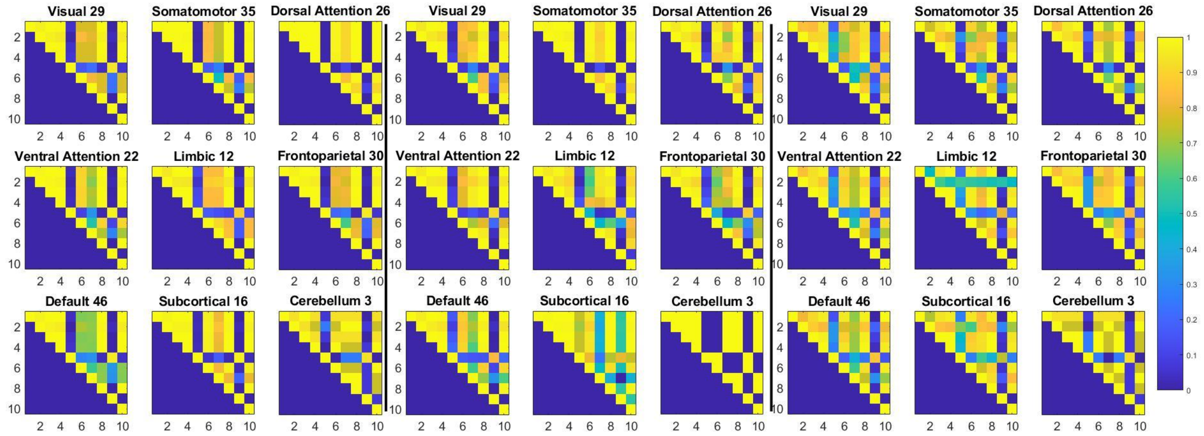


a) Binary structural, b) Binary functional, c) Binary structural weighted by the functional.

**Supplementary Figure 2:** Spearman's correlations computed on average across subjects. Since the matrices are symmetric, only the upper triangular part is shown for visual simplicity. The nine matrices on the left, middle, and right represent the nine RSN's of the SCBIN, FCBIN, and SCFC, respectively. The number next to each RSN indicates the number of nodes from the whole network that belongs to the RSN. Invariant nomenclature; 1 = degree, 2 = betweenness, 3 = closeness, 4 = eigenvector, 5 = clustering, 6 = participation, = 7 within-module degree z-score, 8 = PageRank, 9 =average shortest path, and 10 = subgraph.

**The average distance of a single brain region over all pairs of subjects divided by its standard deviation.**

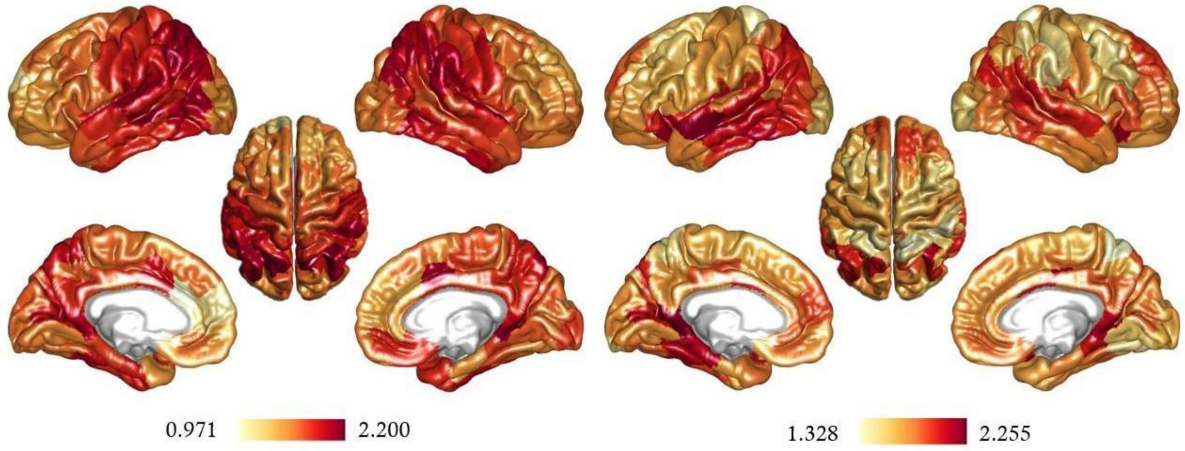


(a) Distance of the structural models (SCBIN on the left and SCWEI on the right).

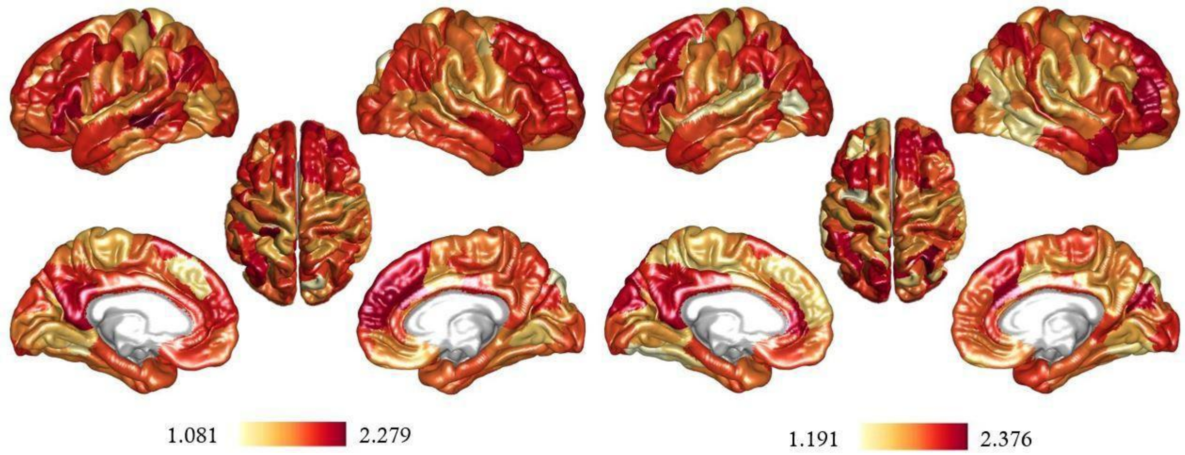


(b) Distance of the functional models (FCBIN on the left and FCWEI on the right).

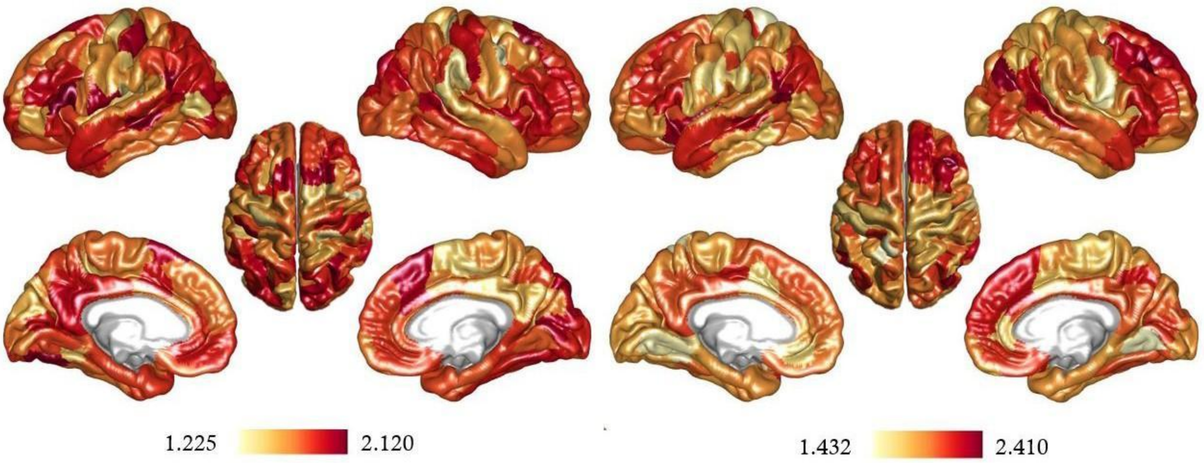


(c) Distance of the SCFC and FCSC models (SCFC on the left and FCSC on the right).

**Supplementary Figure 3:** Multidimensional single region distance. Brain regions exhibiting a high distance across every pair of subjects are in red, and brain regions exhibiting a low distance across every pair of subjects are in yellow. These brain maps represent the ‘graph spatial' network inter-subject variability with red regions being high subject specific brain regions, and white regions being lower subject specific brain regions.

**Global distance between all pairs of subjects at different densities.**

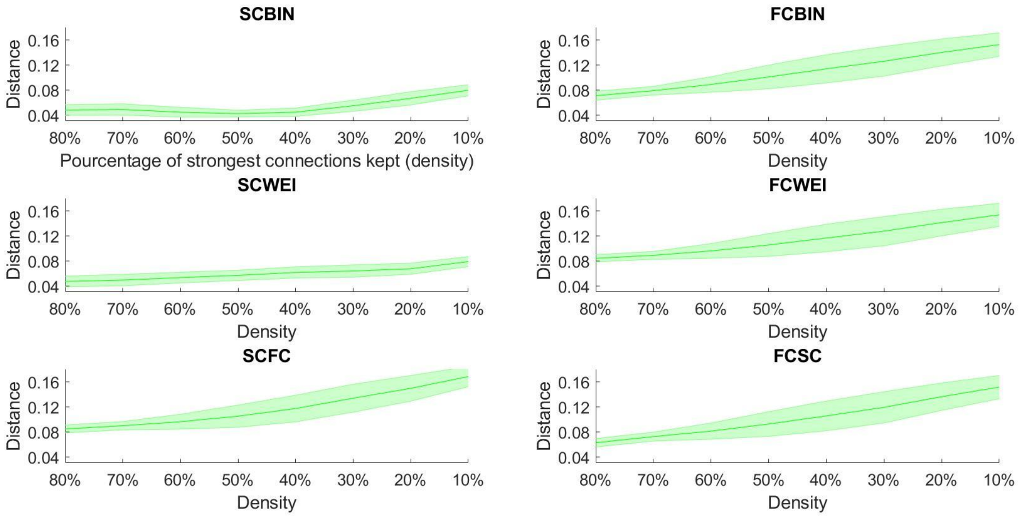


**Supplementary Figure 4:** Multidimensional global distance between connectomes at different densities with shaded areas representing the standard deviation from the mean.

**Global distance between every pair of subjects at each density for every connectome model.**

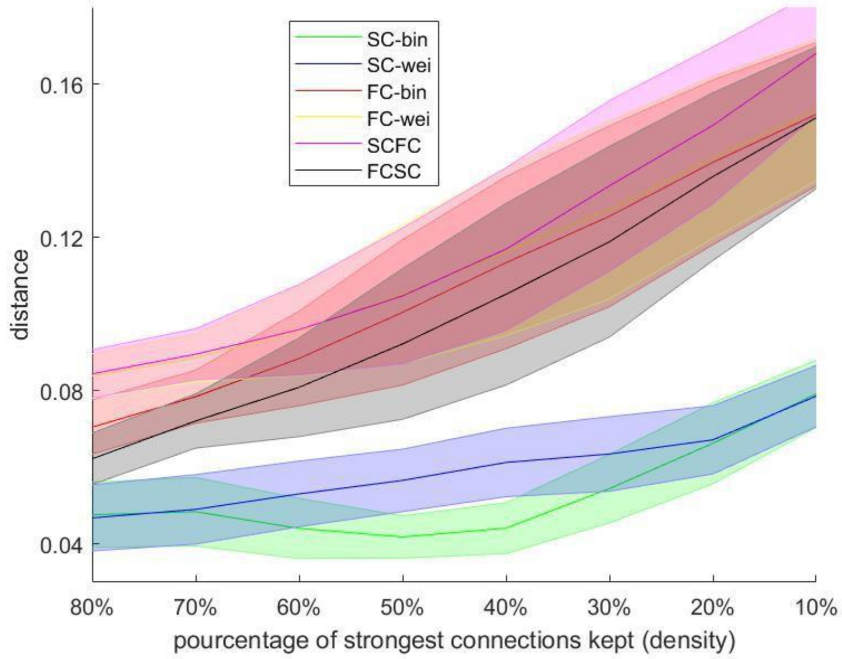


**Supplementary Figure 5:** Multidimensional global distance between connectomes at different densities. All models together can allow direct comparison, for each density, between the models.
